# Diverse novel resident *Wolbachia* strains in Culicine mosquitoes from Madagascar

**DOI:** 10.1101/335786

**Authors:** Claire Louise Jeffries, Luciano M Tantely, Fara Nantenaina Raharimalala, Eliot Hurn, Sébastien Boyer, Thomas Walker

## Abstract

*Wolbachia* endosymbiotic bacteria are widespread throughout insect species and *Wolbachia* transinfected in *Aedes* mosquito species has formed the basis for biocontrol programs as *Wolbachia* strains inhibit arboviral replication and can spread through populations. Resident strains in wild Culicine mosquito populations (the vectors of most arboviruses) requires further investigation given resident strains can also affect arboviral transmission. As Madagascar has a large diversity of both Culicine species and has had recent arboviral outbreaks, an entomology survey was undertaken, in five ecologically diverse sites, to determine the *Wolbachia* prevalence. We detected diverse novel resident *Wolbachia* strains within the *Aedeomyia, Culex, Ficalbia, Mansonia* and *Uranotaenia* genera. *Wolbachia* prevalence rates and strain characterisation through Sanger sequencing with multilocus sequence typing (MLST) and phylogenetic analysis revealed significant diversity and we detected co-infections with the environmentally acquired bacterial endosymbiont *Asaia*. Mosquitoes were screened for major arboviruses to investigate if any evidence could be provided for their potential role in transmission and we report the presence of Rift Valley fever virus in three *Culex* species: *Culex tritaeniorhynchus, Culex antennatus* and *Culex decens*. The implications of the presence of resident *Wolbachia* strains are discussed and how the discovery of novel strains can be utilized for applications in the development of biocontrol strategies.

## Introduction

The endosymbiotic bacterium *Wolbachia* naturally infects approximately 40% of insect species^1^, including mosquito vector species that are responsible for transmission of human diseases. Resident *Wolbachia* strains are present in some major arbovirus disease vectors such as *Culex (Cx.) quinquefasciatus* ^2–4^ and *Aedes (Ae.) albopictus* ^5,6^. *Wolbachia* has been considered for mosquito biocontrol strategies due to the varied phenotypic effects strains have on host mosquito species. Some *Wolbachia* strains can induce a reproductive phenotype termed cytoplasmic incompatibility (CI). This phenotype results in inviable offspring when an uninfected female mates with a *Wolbachia*-infected male. In contrast, *Wolbachia*-infected females produce viable progeny when they mate with both infected and uninfected male mosquitoes. This reproductive advantage over uninfected females allows *Wolbachia* to invade mosquito populations. The CI phenotype was utilized in trials conducted in the late 1960s to eradicate *Cx. quinquefasciatus* from Myanmar ^7^. Releasing large numbers of *Wolbachia-infected* male mosquitoes, to compete with wild type males to induce sterility, is known as the incompatible insect technique (IIT) ^8,9^. Field trials have been undertaken using IIT for both *Ae. albopictus* ^10^ and *Ae. polynesiensis*, a vector of lymphatic filariasis in the South Pacific ^8^.

*Wolbachia* strains have also been shown to protect their native *Drosophila* fruit fly hosts against infection by pathogenic RNA viruses ^11,12^. This discovery led to an alternative biocontrol approach for dengue virus (DENV) transmission which utilizes *Wolbachia*’s ability to inhibit pathogen replication within mosquitoes ^13–17^. *Wolbachia* strains were successfully established in *Ae. aegypti*, the principle mosquito vector of DENV, yellow fever virus (YFV) and Zika virus (ZIKV), through embryo microinjection ^14,18–22^. Preliminary field releases demonstrated the ability of the transinfected wMel strain of *Wolbachia* to establish in wild mosquito populations ^23^ and to provide strong inhibition of DENV in wild mosquitoes ^24^. Further releases are ongoing in DENV endemic countries and mathematical models suggest the wMel strain of *Wolbachia* could reduce the basic reproduction number, R0, of DENV transmission by 66-75% ^25^. *Wolbachia* strains also inhibit other medically important arboviruses including chikungunya virus (CHIKV)^16,26^, YFV ^27^ and recently ZIKV^26,28^.

The prevalence and diversity of natural *Wolbachia* strains in mosquitoes requires further investigation to fully understand how widespread this bacterium is in wild Culicine populations. A recent study using sequence analysis of bacterial 16S rRNA gene amplicons detected a natural strain of *Wolbachia* in *Ae. aegypti* ^29^ suggesting prevalence rates in species previously considered to be uninfected may be underestimated. In *Ae. aegypti* laboratory colonies, the acetic acid bacterium *Asaia* infects the gut and salivary glands and there is competition with *Wolbachia* for colonisation of mosquito reproductive tissues ^30^. *Asaia* has been detected in field populations of *Cx. quinquefasciatus* ^31^ suggesting a complex association between these two endosymbiotic bacteria in wild mosquito populations.

The Culicinae subfamily of mosquitoes (Family Culicidae) contains a large variety of species (>3,000) particularly concentrated in tropical regions. Genera that transmit, or are implicated in transmitting, arboviruses include *Aedeomyia (Ad.), Aedes, Culex, Mansonia (Ma.)* and *Uranotaenia (Ur.)*. Madagascar, a large island located off the southeast coast of mainland Africa, represents an optimal location to investigate *Wolbachia* prevalence in Culicines given its geographic isolation and large diversity of both mosquito species and arboviral diseases. Currently there have been 237 mosquito species morphologically identified in Madagascar in 15 different genera ^32,33^, with 59% of these species thought to be endemic species in Madagascar. There have also been several arbovirus outbreaks in recent history including Rift Valley fever virus (RVFV)^34^, DENV (2006), CHIKV (2006, 2007, 2009 & 2010) and West Nile virus (WNV) ^35^.

Previous studies have identified 64 species in Madagascar that are implicated in transmission of medical or veterinary pathogens ^32^. In this study, we undertook an entomology survey in five ecologically diverse sites in Madagascar to determine the prevalence of resident *Wolbachia* strains in adult female mosquitoes and discovered multiple novel *Wolbachia* strains. The characterisation, typing and phylogeny of these novel *Wolbachia* strains was analysed through Sanger sequencing, with multilocus sequence typing (MLST), to determine the diversity and evolutionary history of resident strains. We also compared *Wolbachia* and *Asaia* prevalence rates and screened mosquitoes for major medically important arboviruses to investigate if there was evidence for the involvement of any of these species as potential arbovirus vectors in these sites.

## RESULTS

### Mosquito abundance and diversity at study sites

Overall, 1174 individuals belonging to seven genera were captured from five collection sites (**Figure 1; Table 1**). *Cx. antennatus* was collected in large numbers from Ambomiharina and Ivato Airport and *Cx. univitattus* was the dominant species in Ambohimarina. 820 (69.85%) of the mosquitoes were caught in CDC light traps, while 354 (30.15%) were caught with Zebu-baited traps. Ambomiharina had the largest number of individuals caught (n=644) and highest diversity, whereas Antafia had the smallest number (n=19), and Ivato Airport the lowest diversity. *Aedes, Aedeomyia* and *Ficalbia (Fi.)* species were collected only at lower elevation plains (Ambomiharina, Antafia) and *Culex* species were collected predominantly at low and high elevation sites (Ambomiharina, Ambohimarina and Ivato airport).

**Figure 1.**
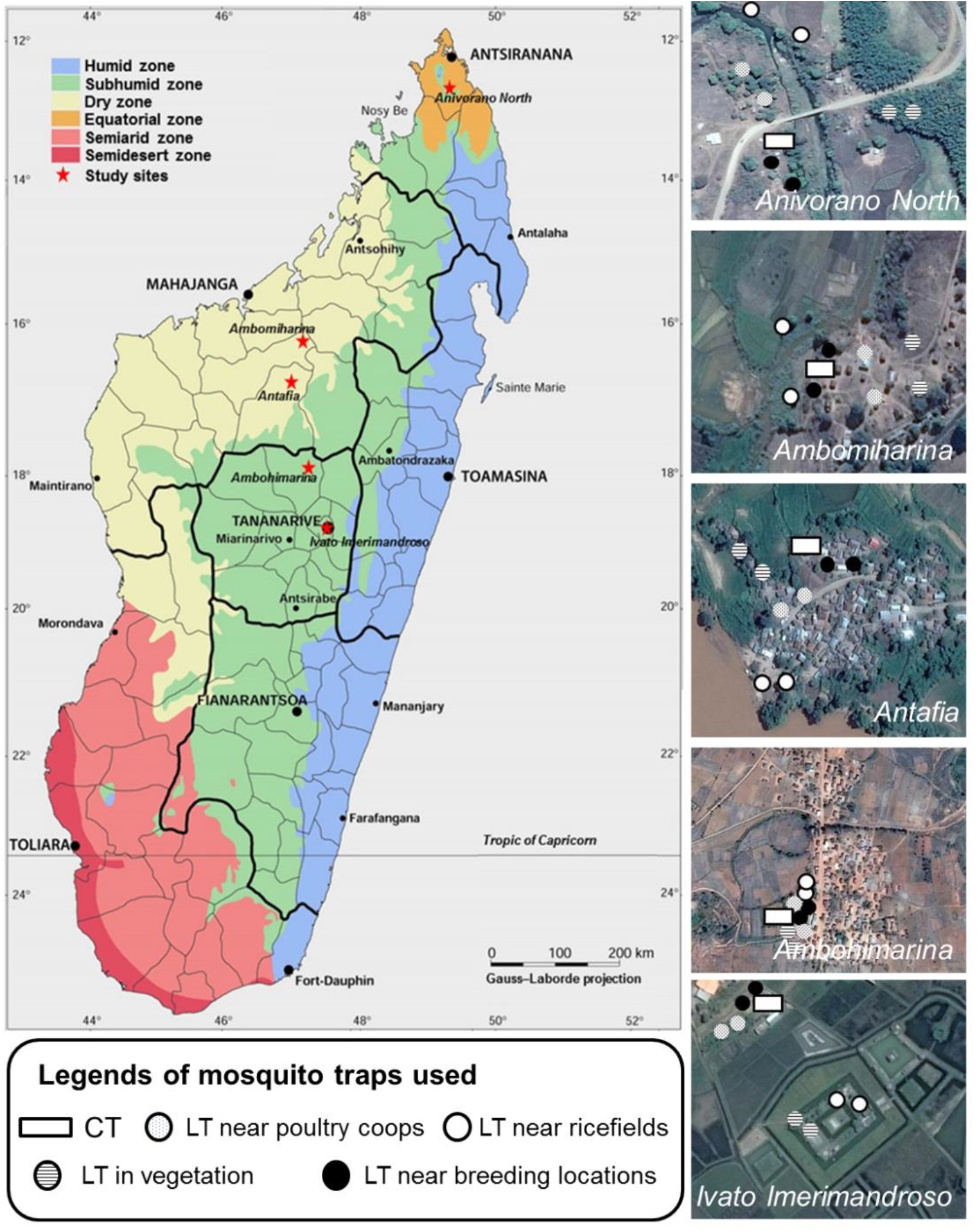
Map of Madagascar showing location of collection sites and mosquito traps used. Adult trapping was undertaken in five ecologically diverse sites using both CDC light traps (LT) and Cattle (zebu) baited traps (CT) near rice fields, poultry coups and breeding locations to maximize the diversity of species collected.

**Table 1.** Summary of mosquito species abundance using morphological identification from five sites in Madagascar. Trapping was undertaken for two consecutive nights with CDC light traps (LT) and cattle (zebu-baited) (CT) traps. Trapping was undertaken between dusk and dawn and mosquitoes from CT traps were collected by mouth aspiration.

|  | Anivorano Nord |  | Ambomiharina |  | Antafia |  | Ambohimarina | Ivato airport |  | Total |
| --- | --- | --- | --- | --- | --- | --- | --- | --- | --- | --- |
|  | CT | LT | CT | LT | CT | LT | LT | CT | LT |  |
| *Aedeomyia furfurea* | 0 | 0 | 0 | 2 | 0 | 0 | 0 | 0 | 0 | 2 |
| *Aedeomyia madagascariensis* | 0 | 0 | 0 | 9 | 0 | 0 | 0 | 0 | 0 | 9 |
| *Aedes aegypti* | 0 | 0 | 0 | 0 | 0 | 2 | 0 | 0 | 0 | 2 |
| *Aedes albocephalus* | 0 | 0 | 0 | 1 | 0 | 0 | 0 | 0 | 0 | 1 |
| *Aedes circumluteolus* | 0 | 0 | 0 | 1 | 0 | 0 | 0 | 0 | 0 | 1 |
| *Culex antennatus* | 8 | 6 | 250 | 91 | 0 | 5 | 52 | 32 | 81 | 525 |
| *Culex bitaeniorhynchus* | 0 | 4 | 1 | 14 | 0 | 1 | 0 | 0 | 0 | 20 |
| *Culex decens* | 0 | 3 | 1 | 12 | 0 | 6 | 0 | 0 | 0 | 22 |
| *Culex giganteus* | 4 | 0 | 0 | 1 | 0 | 0 | 0 | 0 | 1 | 6 |
| *Culex pipiens complex* | 1 | 1 | 0 | 9 | 0 | 0 | 32 | 0 | 1 | 44 |
| *Culex poicilipes* | 0 | 0 | 2 | 17 | 0 | 2 | 1 | 4 | 0 | 26 |
| *Culex tritaeniorhynchus* | 0 | 0 | 25 | 18 | 0 | 0 | 0 | 0 | 0 | 43 |
| *Culex univittatus* | 2 | 6 | 17 | 95 | 0 | 5 | 250 | 0 | 1 | 376 |
| *Ficalbia circumtestacea* | 0 | 0 | 0 | 3 | 0 | 1 | 0 | 0 | 0 | 4 |
| *Lutzia tigripes* | 0 | 1 | 0 | 0 | 0 | 0 | 0 | 0 | 0 | 1 |
| *Mansonia uniformis* | 5 | 10 | 2 | 50 | 0 | 0 | 0 | 0 | 0 | 67 |
| *Uranotaenia spp* | 0 | 5 | 0 | 23 | 0 | 3 | 0 | 0 | 0 | 31 |
| Total | 20 | 36 | 298 | 346 | 0 | 19 | 335 | 36 | 84 | 1174 |

### Detection of resident *Wolbachia* strains and mosquito species identification

We screened individual mosquitoes for the presence of resident *Wolbachia* strains using three conserved *Wolbachia* genes (16 rRNA, wsp and ftsZ) and detected resident strains in six mosquito genera (**Table 2**). The prevalence of *Wolbachia* infection within mosquito species ranged from 3% (1/32) of *Cx. antennatus*, to 100% for *Ad. madagascarica* (9/9). PCR analysis resulted in congruency with the same individual samples amplifying fragments for all three genes, with the exception of *Ma. uniformis, Fi. circumtestacea, Cx. antennatus* and *Cx. duttoni*, in which the wsp gene was not amplified in any individuals. As Madagascar contains diverse mosquito species, we amplified two fragments of the mitochondrial cytochrome *c* oxidase subunit I (CO1) gene ^36,37^ for *Wolbachia-infected* individuals to provide as much molecular confirmation of species as possible given the lack of available CO1 sequences in certain regions of the gene for some species. We were unable to amplify any CO1 gene fragments of *Fi. circumtestacea* but the remaining *Wolbachia*-infected species were confirmed by successful CO1 sequencing (**Figure 2a**). CO1 sequences were deposited in Genbank (accession numbers xxx-yyy) including the first CO1 sequences for *Ad. madagascarica*, and two *Uranotaenia* species from Madagascar, as well as the first *Cx. decens* sequences covering this region of the CO1 gene.

**Table 2.** *Wolbachia* and *Asaia* infection prevalence rates in adult female mosquitoes using three conserved *Wolbachia* genes (*16S rRNA, ftsZ* and *wsp*). An individual was considered *Wolbachia-infected* when any one of the three *Wolbachia* gene fragments were amplified. Species containing resident *Wolbachia* strains are in bold. *Wolbachia* and *Asaia* prevalence rates are shown with 95% confidence intervals in parentheses. Fisher’s exact *post hoc* test P value is shown comparing *Wolbachia* and *Asaia* infections in individual mosquitoes from species that contained at least one individual of both bacterial endosymbionts. ^^^denotes where CO1 sequences were obtained for phylogenetic analysis of mosquito species.

| Mosquito species | number | *Wolbachia* gene PCR amplification |  |  | prevalence (%) |  | Fisher's exact *post hoc* P value<br>(*Wolbachia* vs.<br>*Asaia*) |
| --- | --- | --- | --- | --- | --- | --- | --- |
|  |  | *16S rRNA* | *ftsZ* | *wsp* | *Wolbachia* | *Asaia* |  |
| *Aedes albocephalus* | 1 | 0 | 0 | 0 |  |  |  |
| *Aedes circumlateolus* | 1 | 0 | 0 | 0 |  | 100<br>(20-100) |  |
| *Aedomyia furfuraria* | 2 | 0 | 0 | 0 |  | 100<br>(34-100) |  |
| *Aedomyia madagascarica*<sup>^</sup> | 9 | 9 | 9 | 9 | 100<br>(70-100) | 89<br>(56-98) | >0.99 |
| *Culex antennatus*<sup>^</sup> | 32 | 1 | 1 | 0 | 3<br>(1-16) | 56<br>(38-74) | >0.99 |
| *Culex bitaeniorhynchus* | 12 | 0 | 0 | 0 |  | 58<br>(32-81) |  |
| *Culex decens*<sup>^</sup> | 17 | 3 | 3 | 3 | 18<br>(4-43) | 18<br>(4-43) | 0.46 |
| *Culex duttoni*<sup>^</sup> | 1 | 1 | 1 | 0 | 100<br>(21-100) | 100<br>(21-100) | >0.99 |
| *Culex tritaeniorhynchus*<sup>^</sup> | 19 | 0 | 0 | 0 |  | 95<br>(74-100) |  |
| *Culex giganteus* | 5 | 0 | 0 | 0 |  | 80<br>(37-96) |  |
| *Culex pipiens complex* | 44 | 0 | 0 | 0 |  | 7<br>(1-30) |  |
| *Culex poicilipes* | 21 | 0 | 0 | 0 |  | 48<br>(28-68) |  |
| *Ficalbia circumtestacea* | 3 | 1 | 1 | 0 | 33<br>(6-79) |  |  |
| *Mansonia uniformis*<sup>^</sup> | 24 | 7 | 7 | 0 | 29<br>(15-49) | 67<br>(47-82) | 0.65 |
| *Uranotaenia spp*<sup>^</sup> | 27 | 7 | 7 | 7 | 26<br>(13-45) | 19<br>(8-37) | 0.54 |

**Figure 2.**
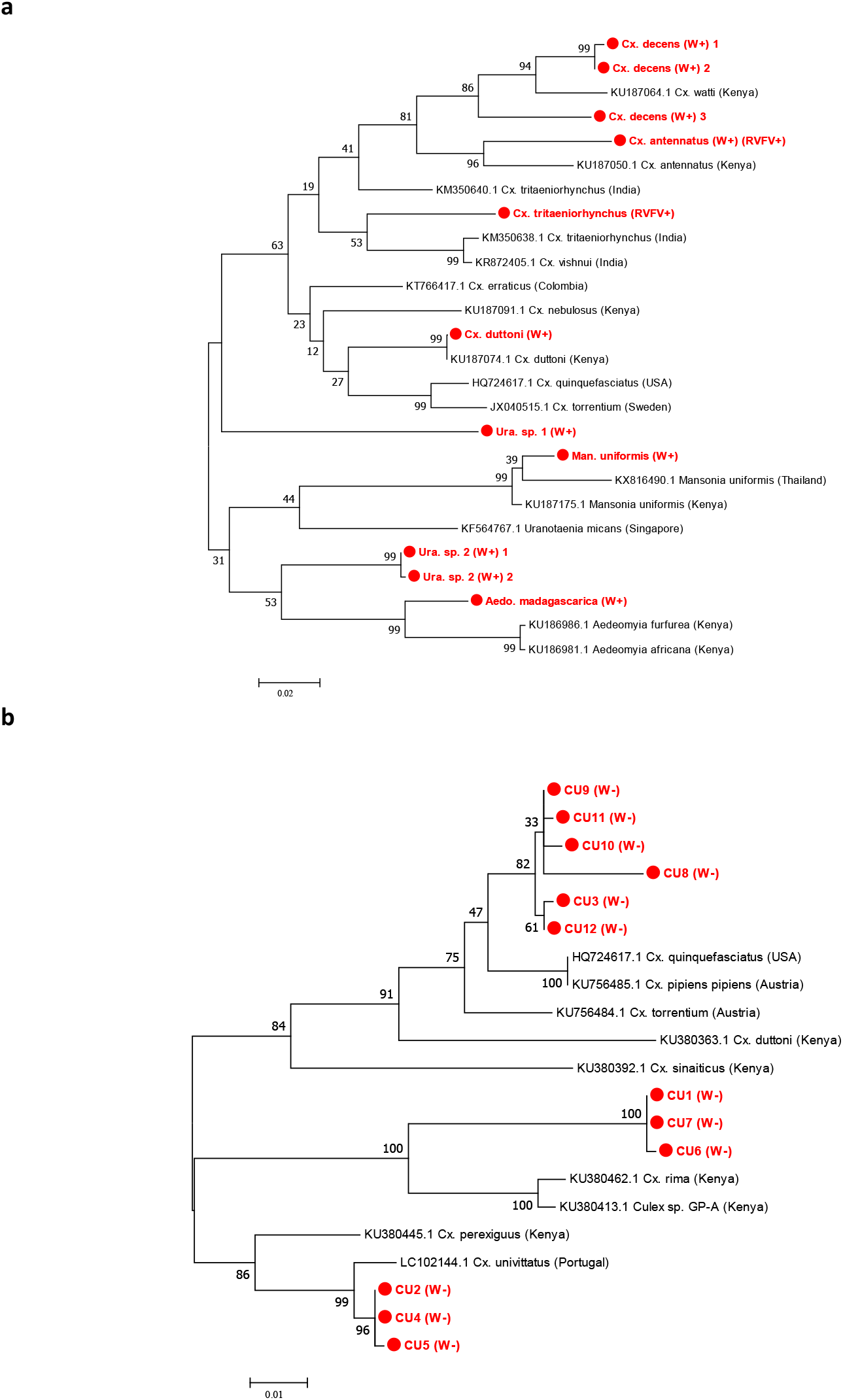
Madagascar mosquito species phylogenetic analysis using the CO1 gene. **a)** Maximum Likelihood phylogenetic analysis of one fragment of the CO1 gene ^37^ showing a tree with the highest log likelihood (−4419.95), comprising 26 nucleotide sequences. There were a total of 669 positions in the final dataset. **b)** Maximum Likelihood phylogenetic analysis for selected *Culex* mosquito species targeting a second CO1 gene fragment ^36^ showing a tree with the highest log likelihood (−2174.09), comprising 21 nucleotide sequences. There were a total of 643 positions in the final dataset. *Wolbachia*-infected = (W+), *Wolbachia*-uninfected = (W-), Rift Valley fever virus-infected =(RVFV+).

Surprisingly, we found no evidence of resident *Wolbachia* strains in members of the *Cx. pipiens* complex indicating the absence of the *w*Pip strain which is usually ubiquitous in *Cx. pipiens pipiens* and *Cx. pipiens quinquefasciatus* sibling species ^38^. As species within the *Cx. pipiens* complex are morphologically indistinguishable, we selected a sub-sample of individuals from different collection sites to determine the species using both species-specific PCR analysis and CO1 sequencing. PCR-based species identification targeting the *ace-2* gene ^39^ revealed the presence of *Cx. pipiens pipiens* individuals in four collection sites (no *Cx. pipiens* complex individuals were collected from Antafia). Confirmation resulted from CO1 phylogenetic analysis through sequencing a second CO1 gene fragment ^36^ and we confirmed six *Wolbachia*-uninfected *Cx. pipiens pipiens* (CU8-CU12) within the *Cx. pipiens* complex (Figure 2b). We also analysed three females (CU1, CU6 and CU7) that were morphologically identified as *Cx. pipiens* complex individuals but did not result in any species confirmation through PCR (**Figure 2b**). CO1 sequencing indicated these individuals were genetically diverse and, with the sequences currently available for comparison, were most closely related to *Cx. rima* and sequences named *Cx. sp.GP-A* from Kenya (94% nucleotide identities, but only 89-94% coverage to the gene fragment amplified), suggesting the possibility of unknown *Culex* species present in Madagascar. We also screened individuals from all mosquito species for *Asaia* and detected the presence of this competing endosymbiont in 13 of 17 species (**Table 2**). Variable prevalence rates were observed and we detected co-infections of *Wolbachia* and *Asaia* in *Ad. madagascarica* (n=8), *Cx. antennatus* (n=1), *Cx. decens* (n=2), *Cx. duttoni* (n=1), *Ma. uniformis* (n=5) and *Uranotaenia* species (n=4). Fisher’s exact *post hoc* tests revealed no association between *Wolbachia* and *Asaia* prevalence rates in species that had at least one individual infected with either bacterial endosymbiont (**Table 2**).

### *Wolbachia* strain characterisations

Sanger sequencing of 16S rRNA, wsp and multilocus sequence typing (MLST) gene fragments was undertaken with *Wolbachia*-infected individuals to characterise, type and determine strain phylogenies. As Madagascar contains some mosquito species that are only present on the island, we predicted the presence of divergent *Wolbachia* strains and this was reflected by variation in the success of amplifying gene target fragments using standard primers and alternative protocols that use M13 sequencing adaptors and / or primers comprising increased degeneracies ^40^. Despite using a variety of primers, we were unable to amplify wsp or any of the five MLST gene loci for *Cx. antennatus* and *Cx. duttoni*. For the remaining resident *Wolbachia* strains, we amplified wsp from all strains except *w*Unif-Mad (**Table 3**). Phylogenetic analysis based on the 16S rRNA gene indicates most of these novel *Wolbachia* strains are clustering with Supergroup B *Wolbachia* strains, such as *w*Pip and *w*Dei (**Figure 3**). Interestingly, only *w*Ura1 clusters with Supergroup A *Wolbachia* strains, such as *w*Mel and *w*Ri. As the wsp gene has been evolving at a faster rate and provides more informative strain phylogenies, with improved phylogenetic resolution ^41^, we compared the phylogenetic relationships for strains that amplified a wsp fragment (**Figure 4**). Similar phylogenetic relationships are shown with wsp, with *w*Ura1 sequences clustering with *w*Mel (Supergroup A), but the remaining strains (*w*Ura2, *w*Dec and *w*Mad) clustered with Supergroup B *Wolbachia* strains.

**Table 3.** Novel resident *Wolbachia* strain wsp and MLST gene allelic profiles. Exact matches to existing alleles present in the database are shown in bold, novel alleles are denoted by the allele number of the closest match and shown in red (number of single nucleotide differences to the closest match). *alternative degenerate primers used to generate sequence. TBA; to be assigned.

| Species | *Wolbachia* strain | WSP |  |  |  |  | MLST alleles |  |  |  |  |  |
| --- | --- | --- | --- | --- | --- | --- | --- | --- | --- | --- | --- | --- |
|  |  | *wsp* | *HVR1* | *HVR2* | *HVR3* | *HVR4* | *gatB* | *CoxA* | *hcpA* | *ftsZ* | *fbpA* | *ST* |
| *Ad. madagascariensis* | wMad | **10** | **10** | **8** | **10** | **8** | **4** | **14** | **40** | **73\*** | **4** | **125** |
| *Uranotaenia spp 1* | wUra1 | 538<br>(2) | **1** | **12** | TBA | **19** | **53** | **50** | 67<br>(3) | 55<br>(1) | **61** | TBA |
| *Uranotaenia spp 2* | wUra2 | 148<br>(4) | **15** | **17** | **87** | **275** | 9<br>(1) | 68<br>(1) | 6<br>(1) | **7** | **10** | TBA |
| *Ma. uniformis* | wUnif-Mad | - | - | - | - | - | **9** | **14** | - | **73** | 4<br>(2) | - |
| *Cx. decens* | wDec | 561<br>(13) | TBA | **232** | **222** | **84** | **9** | 14<br>(2) | 12<br>(1) | 9<br>(2) | 203<br>(1) | TBA |
| *Fi. circumtestacea* | wCir | - | - | - | - | - | 219<br>(2) | 36<br>(2) | 40<br>(1) | 81<br>(1) | **219** | TBA |

**Figure 3.**
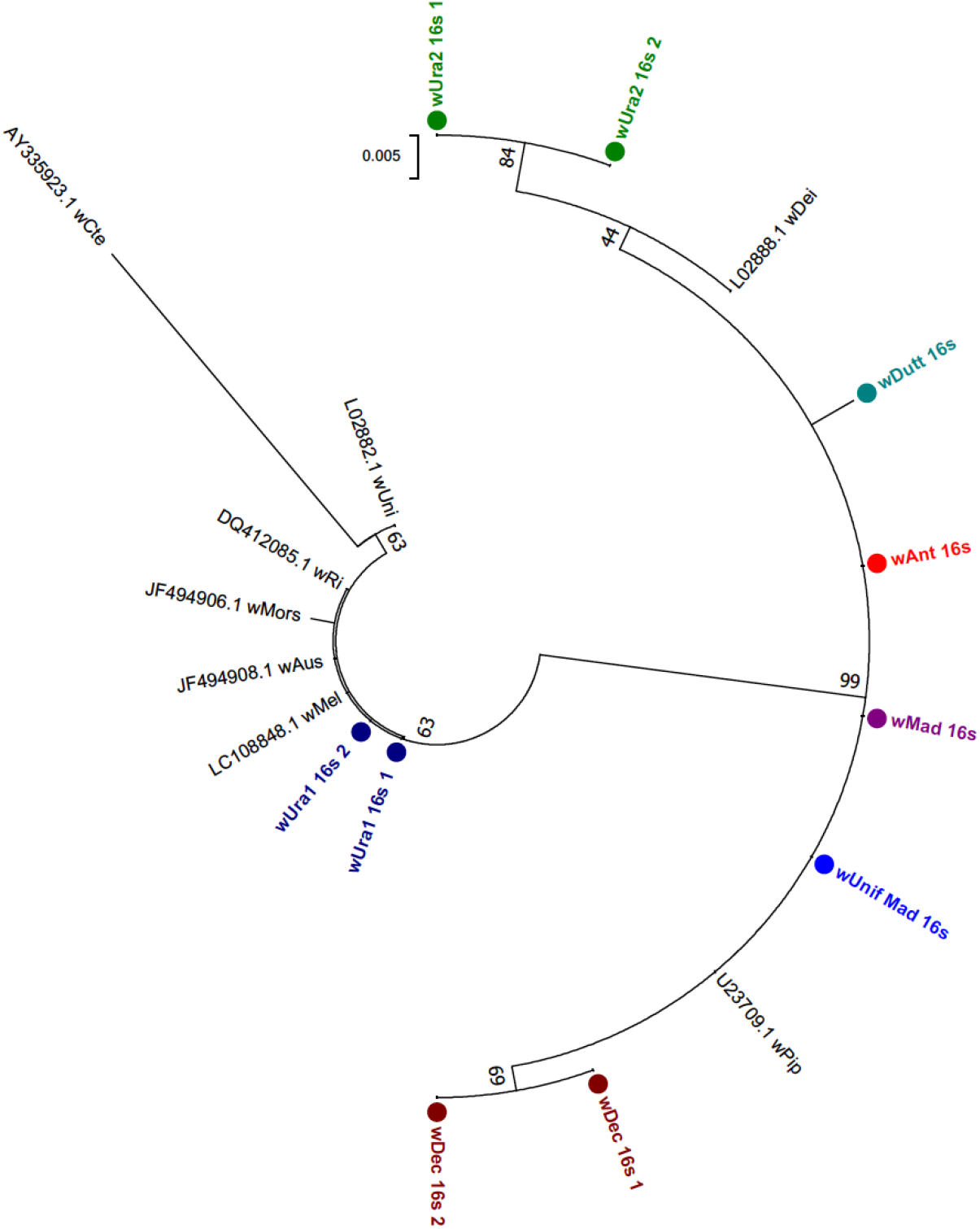
Maximum Likelihood phylogenetic analysis for novel *Wolbachia* strains using the 16S rRNA gene. The tree with the highest log likelihood (−681.38) is shown, comprising 17 nucleotide sequences. There were a total of 335 positions in the final dataset. Resident *Wolbachia* strains in our study are denoted with circles and multiple 16S rRNA sequences from individuals of the same species are included where possible. For comparison, a diverse range of strains available from Genbank were included (shown in black, with accession numbers included).

**Figure 4.**
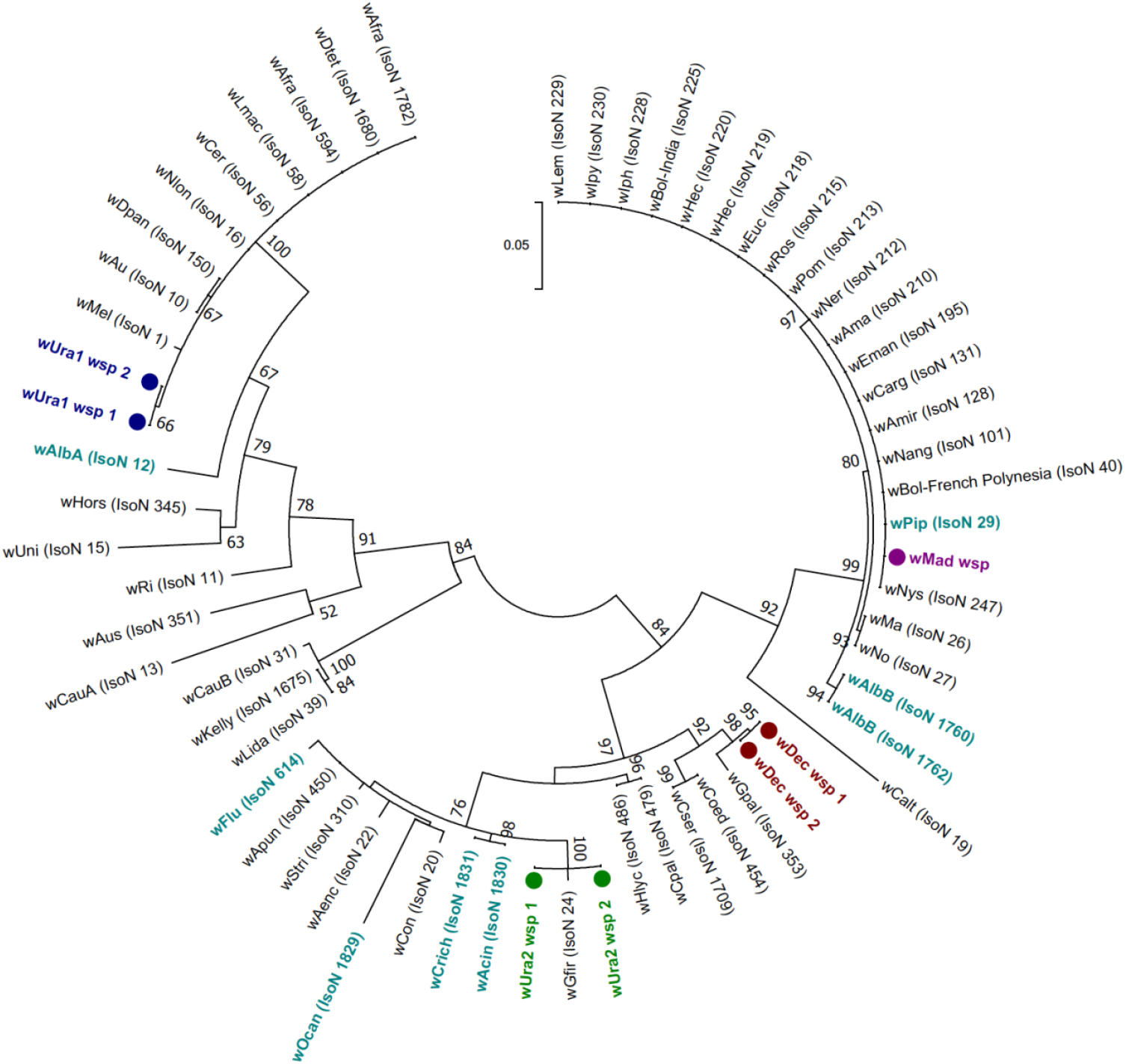
Maximum Likelihood phylogenetic analysis for novel *Wolbachia* strains using the wsp gene. The tree with the highest log likelihood (−2976.84) is shown, incorporating 62 nucleotide sequences. There were a total of 439 positions in the final dataset. Resident *Wolbachia* strains in our study are denoted with circles and multiple wsp sequences from individuals of the same species are included. Wsp sequence data from *Wolbachia* strains downloaded from MLST database for comparison are shown in teal (mosquito strains) and black (all other host strains), with isolate numbers from the MLST database shown in brackets. Novel resident strains discovered and characterised in our study are highlighted in colour and denoted with circles.

MLST resulted in successful sequencing of the majority of gene loci, although we were unable to amplify hcpA from *w*Unif-Mad despite using alternative protocols with degenerate primers (**Table 3**). ftsZ universal primers ^42^ were also needed to obtain a sequence for *w*Mad. Each of the *w*Mad gene sequences produced an exact match to alleles already present in the database, with the resulting allelic profile producing an exact match with strain type 125. This allelic profile matches four isolates currently present in the MLST database, detected in three Lepidopteran species, from French Polynesia, India, Japan and Tanzania (Isolate numbers 40, 247, 270 and 322 respectively), but no isolates from mosquito or other Dipteran hosts. The remaining strains had new alleles for at least one of the MLST gene loci (sequences differed from those currently present in the database by at least one nucleotide difference per locus), resulting in new allelic profiles and therefore new strain types, highlighting the diversity and novelty of these resident strains. All novel allele sequences for each gene locus, complete MLST allelic profiles, and isolate information for each strain, were submitted to the *Wolbachia* MLST database. The phylogeny of these novel strains based on concatenated sequences of all five MLST gene loci (where possible) (**Figure 5**), or four MLST gene loci for analysis of *w*Unif-Mad (hcpA sequence not available) alongside the other strains (**Figure 6**), also reveals significant strain divergence. As expected, due to the general incongruence between host phylogenetic relationships and their resident *Wolbachia* strains ^43,44^, the majority of the novel *Wolbachia* strains in our study provide no evidence for clustering with other strains that infect mosquitoes. One exception is the *w*Mad strain which, although distinct, appears closely related to *w*Pip (the resident strain normally present in the *Cx. pipiens* complex). The other exception is the *w*Unif-Mad strain (based on 4 MLST loci, without hcpA), which is closely related to the *Wolbachia* strain previously detected in *Ma. uniformis* in Kenya (Isolate number 500), and where the pattern of amplification success was also matched as there was absence of amplification for both hcpA and wsp. However, *w*Unif-Mad differs by two nucleotides within the fbpA sequence, producing a novel fbpA allele and therefore a new allelic profile (**Figure 6 and Table 3**).

**Figure 5.**
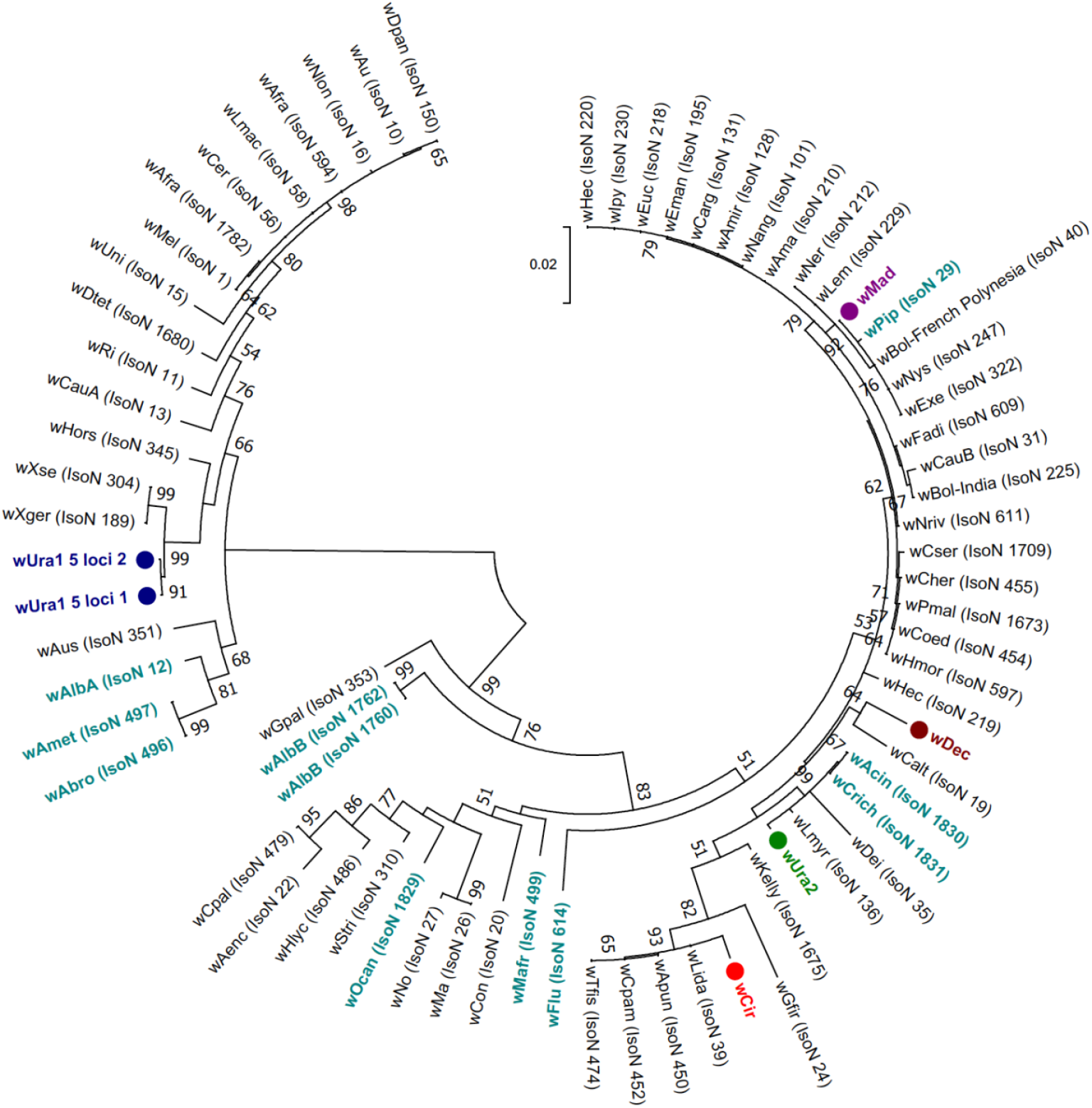
Maximum Likelihood molecular phylogenetic analysis from concatenation of the five MLST loci. The tree with the highest log likelihood (−9010.45) is shown incorporating 73 nucleotide sequences. There were a total of 2063 positions in the final dataset. Concatenated sequence data from *Wolbachia* strains downloaded from the MLST database for comparison are shown in teal (mosquito strains) and black (all other host strains), with isolate numbers from the MLST database shown in brackets. Novel resident strains discovered and characterised in our study are highlighted in colour and denoted with circles.

**Figure 6.**
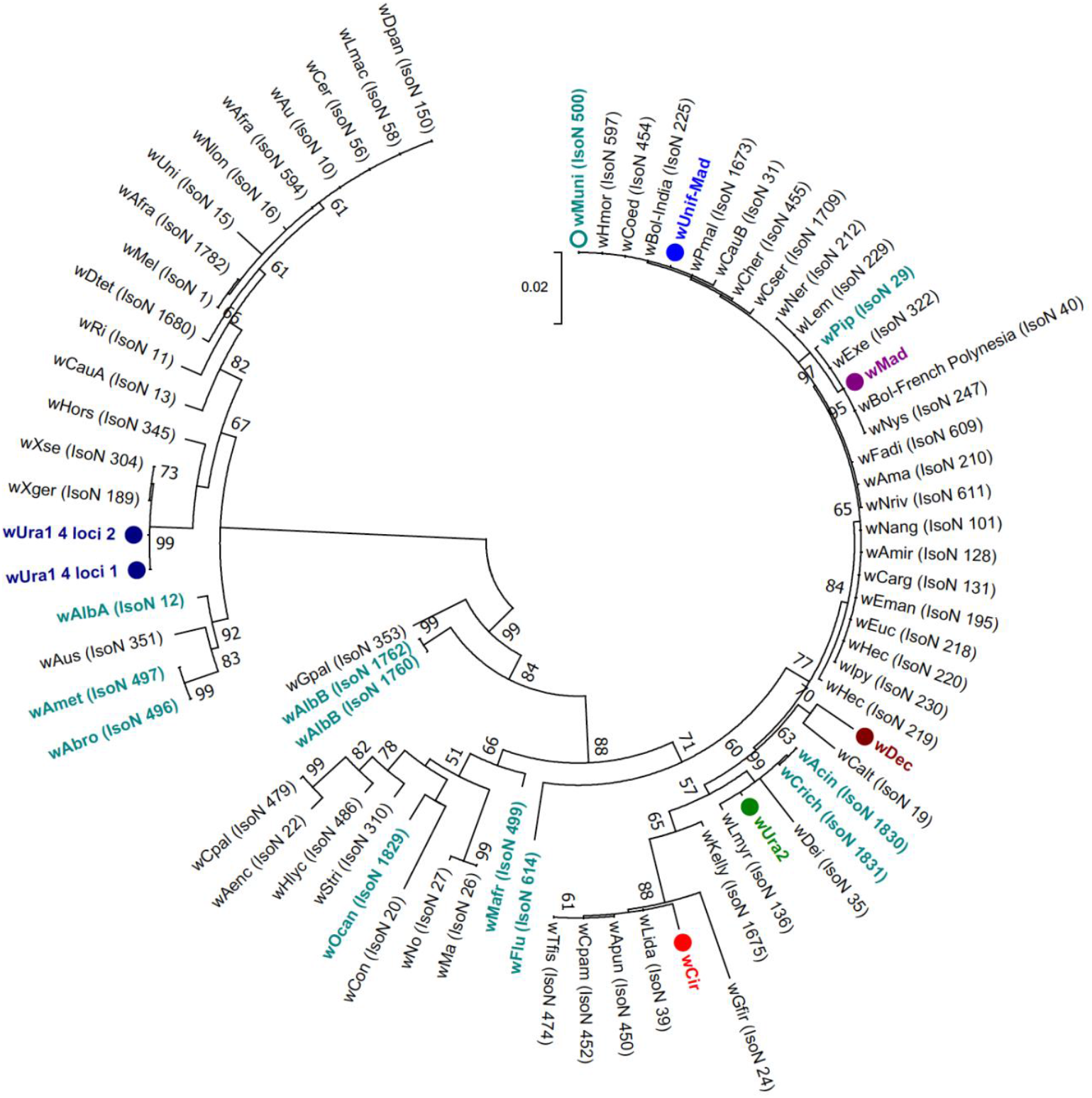
Maximum Likelihood phylogenetic analysis from concatenation of four MLST loci (coxA, fbpA, ftsZ and gatB). The tree with the highest log likelihood (−7114.64) is shown incorporating 75 nucleotide sequences. There were a total of 1624 positions in the final dataset. Concatenated sequence data from *Wolbachia* strains downloaded from the MLST database for comparison are shown in teal (mosquito strains) and black (all other host strains), with isolate numbers from the MLST database shown in brackets. Concatenated *Ma. uniformis* sequence data from Kenya ^51^ (Isolate number 500) downloaded from the MLST database is highlighted in teal and denoted with a clear circle, for comparison with the *w*Unif-Mad strain (blue with filled circle) detected in this study. Other novel resident strains discovered and characterised in our study are highlighted in colour and denoted with filled circles.

### Site locations for *Wolbachia*-infected mosquitoes

The novel resident *Wolbachia* strain in *Cx. decens*, named *w*Dec, was only found in individuals collected from Ambomiharina (West domain) with no detectable infections in individuals from Anivorano North (North domain) or Antafia (West domain). Resident *Wolbachia* strains in both *Cx. antennatus* (*w*Ant) and *Cx. duttoni* (*w*Dut) were from individuals collected in Ambomiharina (West domain). The novel *w*Mad *Wolbachia* strain was detected in all *Ad. madagascarica* females collected in Ambomiharina (West domain). We detected the *w*Unif-Mad strain in *Ma. uniformis* from both Ambomiharina (West domain) and Anivorano North (North domain). Our single *Wolbachia-infected Fi. circumtestacea* (*w*Cir) was collected from Ambomiharina (West domain) and novel *Wolbachia* strains in *Uranotaenia* species (*w*Ura1 and *w*Ura2) were detected in individuals from Anivorano Nord (located in the Northern domain), Ambomiharina and Antafia (Western domain).

### Arbovirus detection in mosquitoes

In total, 704 mosquitoes of species considered potential vectors of medically important arboviruses in Madagascar were screened using arbovirus-specific PCR assays. Mosquitoes were screened as either individuals or pools of 3-5 individuals from the same location and trapping method (**Table 1**). RVFV RNA was detected in two individually screened female adults of *Cx. decens*: one collected from Ambomiharina and one from Antafia. We also detected RVFV in ten individual *Cx. tritaeniorhynchus* females and five individual *Cx. antennatus* females from Ambomiharina. We sequenced RVFV PCR products from all positive individuals to confirm correct target amplification and also used CO1 sequencing to confirm mosquito species (representative samples are shown in **Figure 2a**). No evidence of infection for the remaining arboviruses was present from any mosquitoes collected from the five sites in our study.

## DISCUSSION

Madagascar contains some of the most diverse and unique flora and fauna given its geographical isolation. There are currently 237 known species of mosquitoes present on the island, but recent newly described species ^32^ would suggest the possibility of more species present only in Madagascar. In our study, we assessed the *Wolbachia* prevalence of mosquito species that have the potential to be arbovirus vectors from five diverse ecological sites. To our knowledge, we are reporting for the first time resident *Wolbachia* strains in seven mosquito species: *Ad. madagascarica, Cx. antennatus, Cx. decens, Cx. duttoni, Fi. circumtestacea* and two *Uranotaenia* species. The novel resident *Wolbachia* strain in *Cx. decens*, named *w*Dec, was only found in individuals collected from one location (Ambomiharina in the West domain) and an overall prevalence of 20% indicates that prevalence rates are likely to be variable across the island. *Cx. decens* is present in all biogeographic domains of Madagascar and is particularly abundant in the central domain ^45^. In Madagascar, this species was previously shown to be infected with WNV and Babahoyo virus ^45^ and in mainland Africa has been shown to be infected with additional arboviruses including Moussa virus ^46^. The wMad strain was found in all *Ad. madagascarica* individuals we collected from Ambomiharina (West domain). *Ad. madagascarica* was previously found naturally infected with WNV and is zoophilic, with a preference for feeding on avian blood ^47^, demonstrating its likely involvement in zoonotic arbovirus transmission in Madagascar. In previous entomological surveys in north-western Madagascar where WNV is endemic, *Ad. madagascarica* was the most abundant species and WNV RNA was detected in one pool suggesting this species may play a role in WNV maintenance or transmission ^48^.

Novel resident *Wolbachia* strains were also detected in two *Uranotaenia* species, of which there is very limited knowledge of these species in Madagascar. The lack of available CO1 sequences prevented molecular species confirmation but *Ur. anopheloides* is endemic to Madagascar and the Comoros archipelago and is most abundant is warmer regions of the western domain ^32^. *Ur. alboabdominalis* is thought to occur mainly in the eastern and western domains of Madagascar and *Ur. neireti* in the central and eastern domains between 900-200m above sea level. There is also limited knowledge of *Fi. circumtestacea* in Madagascar, with reports of its presence in the eastern and western domains ^32^, but this species is not currently confirmed as being involved in arbovirus transmission. As we were only able to amplify a fragment of the 16S rRNA gene for resident strains in *Cx. antennatus* (*w*Ant) and *Cx. duttoni* (*w*Dut), further studies are needed to determine if the lack of additional *Wolbachia* gene amplification is due to strain variability or low-density infections preventing successful amplification.

*Ma. uniformis* is present in all biogeographical domains of Madagascar, is considered an anthropophilic vector of numerous arboviruses such as RVFV ^49^ and WNV ^47^, and is a known vector of *Wuchereria bancrofti* filarial nematodes in numerous countries. Resident *Wolbachia* strains in *Ma. uniformis* have previously been shown in southeast Asia ^50^, Kenya ^51^ and more recently in Sri Lanka ^52^. Interestingly in Sri Lanka, 3/3 (100%) of *Ma. uniformis* individuals amplified the wsp gene ^52^ but in our study all seven individuals failed to amplify wsp. *Ma. uniformis* was also one of the four *Wolbachia-infected* mosquito species from populations in Kenya ^51^ and this provides the only comparative MLST and reports the requirement for nested PCR to amplify hcpA. Our *Wolbachia* prevalence rate of 29% (7/24) for *Ma. uniformis* is similar to the 29% (5/19) reported in Kenyan populations ^51^ suggesting this species has variable prevalence rates in wild populations.

The absence of *Wolbachia* infections in the *Cx. pipiens* complex (particularly *Cx. pipiens pipiens* which were confirmed by PCR and CO1 sequencing) was surprising given high infection rates of the wPip strain are often observed in wild populations ^53–57^. Furthermore, our results contrast with a study undertaken in Madagascar as part of a wider Incompatible Insect Technique (IIT) from southwestern Indian Ocean islands in which the wPip strain was detected in *Cx. pipiens quinquefasciatus* mosquitoes from Antananarivo ^58^. This suggests the possibility of variable infection rates between *Cx. pipiens pipiens* and its sibling species *Cx pipiens quinquefasciatus* in Madagascar populations (which we did not collect in our study). We also undertook molecular identification of three *Culex* individuals (also Wolbachia-uninfected) that were phylogenetically similar to *Cx. rima* (**Figure 2b**) which is consistent with a previous entomological survey that describes an unknown species morphologically similar to *Cx. rima* ^59^. Further phylogenetic studies are warranted to determine the diversity of *Culex* species and the variability of *Wolbachia* prevalence within the *Cx. pipiens* complex.

The presence of resident *Wolbachia* strains in mosquito vector species which can transmit human arboviruses could be influencing arboviral transmission dynamics in field populations of Madagascar. In laboratory studies, resident *Wolbachia* strains have been shown to impact arboviral transmission. For example, *Wolbachia* was shown to reduce DENV infection of salivary glands and limit transmission in *Ae. albopictus* ^60^. Outbreaks of RVFV in Madagascar are thought to result from infected domestic animals imported from east Africa ^61^ and *Cx. antennatus* has previously been identified as an important RVFV vector in Madagascar ^34^. Our results would indicate *Cx. tritaeniorhynchus* is also likely contributing to transmission in Ambomiharina given we also detected RVFV RNA in multiple non-blood fed females of this species. RVFV has not previously been detected in this species in Madagascar, however, *Cx. tritaeniorhynchus* was implicated as a major vector during a large outbreak of RVFV in Saudi Arabia ^62^ so this species contribution to transmission in Madagascar could potentially be currently underestimated. The detection of RVFV in three *Culex* species at Ambomiharina and *Cx. decens* at Antafia would correlate with previous studies that have shown that RVFV is one of the most abundant and widely distributed arboviruses across the island and there have been several recent RVFV outbreaks ^34,63^. In our study, we detected RVFV in Wolbachia-uninfected *Cx. decens* and *Cx. tritaeniorhynchus* individuals but our single *Wolbachia-infected Cx. antennatus* was co-infected with RVFV. This female was non-blood fed indicating potential dissemination of RVFV beyond the bloodmeal. We were also unable to amplify any *Wolbachia* genes other than 16S rRNA from this individual which could be explained by a very low-density resident strain present in *Cx. antennatus. Wolbachia* tissue tropism influences the potential to inhibit arboviruses ^15,64^ as some resident *Wolbachia* strains are present predominantly in reproductive tissue and have little effects on arboviruses ^65^. A low density wAnt infection may also explain why we only detected *Wolbachia* infections in 1 of 32 *Cx. antennatus* females in our study. However, as arbovirus infection rates are normally low in mosquito populations, correlating the prevalence of *Wolbachia* and RVFV would require a significantly larger number of mosquitoes.

The co-detection of resident *Wolbachia* strains and *Asaia* in seven species also indicates the potential for complex tripartite interaction with arboviruses. In our study, *Asaia* was also detected in the single *Wolbachia*-infected *Cx. antennatus* individual so further studies should be undertaken to determine the dynamics of these microbes within wild mosquito populations. *Asaia* has previously been detected in field-caught *Ae. albopictus* from Madagascar with a prevalence rate of 46% ^66^ but *Asaia* was one of 27 bacterial genera present in populations sampled from different Madagascar biotypes ^67^ highlighting the complexity of the microbiota. A similar study analysing *Ae. aegypti* and *Ae. albopictus* populations in Madagascar (including resident *Wolbachia* strains) concluded that the environment was a factor in the overall microbiota composition and diversity ^68^. It has also recently been shown that there is no evidence for *Wolbachia-Asaia* co-infections in a wide range of *Anopheles* species and the environment likely influences *Asaia* prevalence and density in wild mosquito populations ^69^.

The discovery of novel resident *Wolbachia* strains in mosquito species from Madagascar may also impact future attempts to extend *Wolbachia* biocontrol strategies by using these strains for applied use through transinfection. The naturally occurring wAlbA and wAlbB strains of *Wolbachia* that infect *Ae. albopictus* have been successfully transferred to *Ae. aegypti* ^14,18,21^ and significantly inhibit DENV ^14,21^. This suggests the introduction of novel resident *Wolbachia* strains from other Culicine species can generate inhibitory effects on arboviruses. Further experiments should elucidate both the density and tissue tropism of these novel strains to determine candidate strains for transfer to species such as *Cx. quinquefasciatus* that are major arbovirus vectors. As *Cx. quinquefasciatus* contain a resident *Wolbachia* strain (wPip) ^70^, introduction of ‘transinfected’ strains would require a stable association, with introduced strains growing to higher densities in specific tissues which result in inhibition of pathogen transmission ^64^. Our discovery of novel resident strains in *Culex* species closely related to *Cx. quinquefasciatus* may improve transinfection success and ultimately lead to *Wolbachia* biocontrol strategies targeting *Culex* species.

## METHODS

### Study sites

Five study sites were chosen according to a large variety of potential ecological factors in order to sample a wide variety of mosquito species (**Figure 1**). Anivorano Nord (located in the Northern domain), Ambomiharina and Antafia (Western domain), Ambohimarina and Ivato Imerimandroso (Central domain) were sites selected to sample mosquito populations. Anivorano North (12°45’52.2”S, 49°14’19.3”E, at 375 m above sea level (asl)) belongs to the district of Antsiranana II and is located 75km south of the town of Antsiranana. Our study was performed in a village 500 m south of the city of Anivorano Nord. The village consists of 32 houses distributed on both sides of the Irodo River. This river irrigates rice fields with heterogeneous rice phenology that are restricted to a small valley floor. Ambomiharina (16°21’62.2”S, 46°59’34.4”E, at 84 m asl) is located in the Ambato-Boeny district within the Tsaramandroso municipality. The village consists of 35 houses and is located 5 km south east of Ankarafantsika Forest, surrounded by a large rice field, irrigated by numerous large streams. Antafia (17°01’37.8”S, 46°45’61.7”E, at 64 m asl) is located on the Betsiboka River, at 11 km south west of the town of Maevatanana and consists of 193 houses. The landscape consists of Betsiboka River in the west and large dry rice fields in the south, east and north of the village. Additionally, this site is characterized by a resident population of bats that were found both in houses and flying during dusk. Ambohimarina (18°19’58.6”S, 47° ‘06.33.3”E, at 1212 m asl) belongs to the Ankazaobe district, 5 km south of the city of Ankazobe. The village consists of 289 houses, is located on a hill and surrounded with valleys composed of rice fields. Ivato Imerimandroso (18°47’31.6”S, 47°28’68.4”E, at 1261 m asl) is located 300 m from Ivato Airport in the city of Antananarivo. Rice fields are flooded and surrounded with swamp and marshes. The domestic animals and livestock consist of cattle, dogs, and poultry in all five study sites.

### Mosquito sampling

Mosquito sampling was undertaken utilizing both CDC light traps and a net trap baited with Zebu (local species of cattle) to attract zoophilic species at each of the five study sites ^63^. Trapping was performed over two consecutive nights per site, with CDC light traps set up at dusk to be operational during sunset and overnight, before being collected at dawn. Eight CDC light traps were placed around each site nearby dwellings, near poultry coops, inside areas of denser vegetation (typically forested areas or crop fields) and near potential breeding locations which varied depending on each site. At each site, one Zebu trap was placed in a way which simulated local animal husbandry practices (i.e. inside of a shed, nearby to a large corral, or on its own tied to a post). Zebu traps were set up following sunset and mosquitoes were collected by mouth aspiration before dawn.

### Morphological identification and RNA extraction

Mosquitoes were anesthetized with chloroform vapor and morphological identification was performed using a variety of mosquito genera keys ^32^. Following identification, mosquitoes were placed into 96 well storage plates or Eppendorf tubes and RNAlater (Sigma, UK) was added prior to storage at 4°C or lower to prevent viral RNA degradation. RNA was extracted from individual whole mosquitoes (or additional pools for further detection of human pathogens) using QIAGEN RNeasy 96 kits according to manufacturer’s instructions. RNA was eluted in 40 μL of RNase-free water and stored at −80°C. A QIAGEN QuantiTect Reverse Transcription kit was used to reverse transcribe RNA, generating cDNA from all RNA extracts, according to manufacturer’s instructions.

### Molecular species identification

Selected samples morphologically identified as part of the *Cx. pipiens* complex were further analyzed using multiplex PCR targeting the ace-2 locus ^39^ and Sanger sequencing of the mitochondrial gene cytochrome oxidase I gene (CO1), shown to provide the optimal discrimination of *Culex pipiens* complex species ^36,37^.

### *Wolbachia* screening

*Wolbachia* detection was first undertaken targeting three conserved *Wolbachia* genes previously shown to amplify a wide diversity of strains; 16S rDNA gene ^71^, *Wolbachia* surface protein (wsp) gene ^41^ and FtsZ cell cycle gene ^72^. DNA extracted from a *Drosophila melanogaster* fruit fly (infected with the wMel strain of *Wolbachia*) was used a positive control, in addition to no template controls (NTCs). 16S rDNA and wsp gene PCR reactions were carried out in a Bio-Rad T100 Thermal Cycler using standard cycling conditions ^41,71^ and PCR products were separated and visualised using 2% E-gel EX agarose gels (Invitrogen) with SYBR safe and an Invitrogen E-gel iBase Real-Time Transilluminator. FtsZ gene real time PCR reactions were prepared using 5μl of FastStart SYBR Green Master mix (Roche Diagnostics), a final concentration of 1μM of each primer, 1μL of PCR grade water and 2μL template DNA, to a final reaction volume of 10μL. Prepared reactions were run on a Roche LightCycler® 96 System for 15 minutes at 95°C, followed by 40 cycles of 95°C for 15 seconds and 58°C for 30 seconds. Amplification was followed by a dissociation curve (95°C for 10 seconds, 65°C for 60 seconds and 97°C for 1 second) to ensure the correct target sequence was being amplified. PCR results were analysed using the LightCycler® 96 software (Roche Diagnostics). *Asaia* detection was also undertaken targeting the 16S rRNA gene ^73,74^.

### *Wolbachia* MLST

Multilocus sequence typing (MLST) was undertaken to characterize *Wolbachia* strains using the sequences of five conserved genes as molecular markers to genotype each strain. In brief, 450-500 base pair fragments of the coxA, fbpA, hcpA, gatB and ftsZ *Wolbachia* genes were amplified from one individual from each *Wolbachia-infected* mosquito species using previously optimised protocols ^40^. A *Culex pipiens* gDNA extraction (previously shown to be infected with the wPip strain of *Wolbachia*) was used a positive control for each PCR run, in addition to no template controls (NTCs). If no amplification was detected using standard primers, further PCR analysis was undertaken using alternative protocols which included utilisation of M13 sequencing adaptors and / or primers with increased degeneracies ^40^.

### Sanger sequencing

PCR products were separated and visualised using 2% E-gel EX agarose gels (Invitrogen) with SYBR safe and an Invitrogen E-gel iBase Real-Time Transilluminator. PCR products were submitted to Source BioScience (Source BioScience Plc, Nottingham, UK) for PCR reaction clean-up, followed by Sanger sequencing to generate both forward and reverse reads. Sequencing analysis was carried out in MEGA7 ^75^ as follows. Both chromatograms (forward and reverse traces) from each sample was manually checked, analysed, and edited as required, followed by alignment by ClustalW and checking to produce consensus sequences. Consensus sequences were used to perform nucleotide BLAST (NCBI) database queries, and *Wolbachia* gene loci sequences were used for searches against the *Wolbachia* MLST database (http://pubmlst.org/wolbachia) ^76^. If a sequence produced an exact match in the MLST database, we assigned the appropriate allele number, otherwise new alleles were added and complete MLST profiles submitted to the *Wolbachia* MLST database.

### Phylogenetic analysis

Maximum Likelihood phylogenetic trees were constructed from Sanger sequences as follows. The evolutionary history was inferred by using the Maximum Likelihood method based on the Tamura-Nei model ^77^. The tree with the highest log likelihood in each case is shown. The percentage of trees in which the associated taxa clustered together is shown next to the branches. Initial tree(s) for the heuristic search were obtained automatically by applying Neighbor-Join and BioNJ algorithms to a matrix of pairwise distances estimated using the Maximum Composite Likelihood (MCL) approach, and then selecting the topology with superior log likelihood value. The trees are drawn to scale, with branch lengths measured in the number of substitutions per site. Codon positions included were 1st+2nd+3rd+Noncoding. All positions containing gaps and missing data were eliminated. The phylogeny test was by Bootstrap method with 1000 replications. Evolutionary analyses were conducted in MEGA7 ^75^.

### Arbovirus screening

Arbovirus screening using published PCR assays included the major arboviruses of public health importance, suspected or having the potential of being transmitted in Madagascar (supplementary Table 1). PCR reactions for all assays except ZIKV were prepared using 5 μL of Qiagen SYBR Green Master mix, a final concentration of 1 μM of each primer, 1 μL of PCR grade water and 2 μL template cDNA, to a final reaction volume of 10 μL. Prepared reactions were run on a Roche LightCycler® 96 System and PCR cycling conditions are described in supplementary Table S1. Amplification was followed by a dissociation curve (95°C for 10 seconds, 65°C for 60 seconds and 97°C for 1 second) to ensure the correct target sequence was being amplified. Zika virus screening was undertaken using a Taqman probe-based assay using 5 μL of Qiagen QuantiTect probe master mix, a final concentration of 1 μM of each primer, 1 μL of PCR grade water and 2 μL template cDNA, to a final reaction volume of 10 μL. PCR results were analysed using the LightCycler^®^ 96 software (Roche Diagnostics). Synthetic long oligonucleotide standards (Integrated DNA Technologies) of the amplified PCR product were generated in the absence of biological virus cDNA positive controls and each assay included negative (no template) controls.

### Statistical analysis

Fisher’s exact *post hoc* test in Graphpad Prism 6 was used to compare *Wolbachia* and *Asaia* prevalence rates in species that had at least one individual infected with *Wolbachia*.

#### Ethics approval and consent to participate

Ethical clearance for the use of Zebu cattle in baited traps was obtained from the ethical committee of the Livestock ministry of Madagascar which is the sole relevant authority for animal care in Madagascar. The ethical committee number is: 2012/WN/Minel/3. We followed the European guidelines (European directives EU 86/609-STE123 and 2010/63/EU) for animal handling. To minimize the risk of infection of mosquito borne diseases, an internal net was used to protect the cattle against mosquito bites.

## Availability of data and material

All data generated or analysed during this study are included in this article.

## Acknowledgements

We would like to thank all the villagers who participated in the fieldwork. Funding was provided by the London School of Hygiene and Tropical Medicine, Bayer Pharmaceuticals and a Wellcome Trust /Royal Society grant awarded to TW (101285/Z/13/Z): http://www.wellcome.ac.uk; https://royalsociety.org. The funders had no role in study design, data collection and analysis, decision to publish, or preparation of the manuscript.

## Author contributions statement

All authors contributed to the design of the study. SB facilitated the field study that was carried out by LMT, FNR and EH involving mosquito capture and morphological identification in the field. CLJ, EH and TW performed molecular analysis of samples. CLJ performed Sanger sequencing analysis, MLST and phylogenetic analysis. All authors read and approved final version of the manuscript.

---

**Supplementary table S1.**
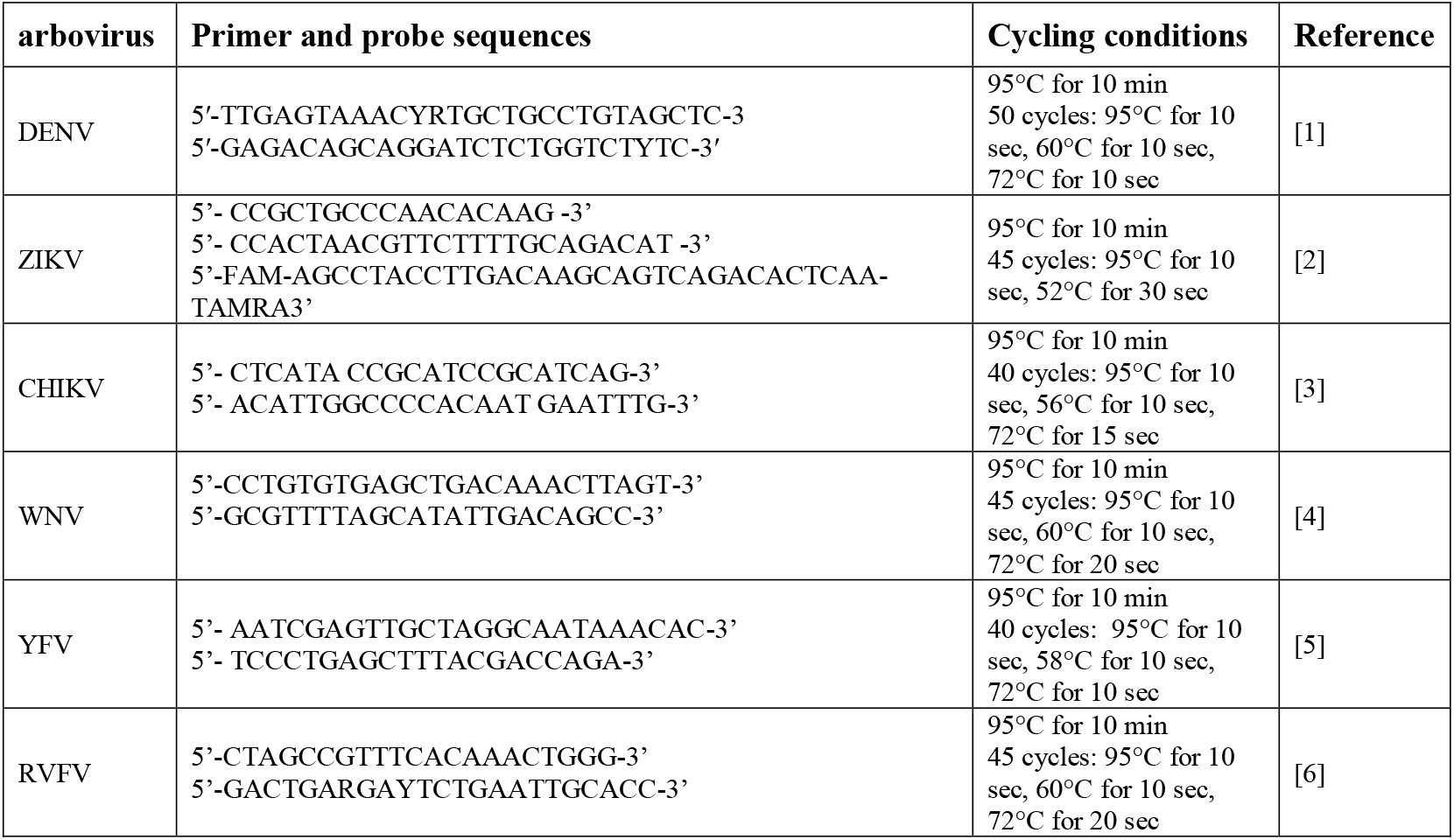
Arbovirus screening assays including PCR primer/probes sequences and cycling conditions used to screen mosquitoes.

